# Neutralization Escape by SARS-CoV-2 Omicron Subvariant BA.2.86

**DOI:** 10.1101/2023.09.04.556272

**Authors:** Ninaad Lasrado, Ai-ris Y. Collier, Nicole P. Hachmann, Jessica Miller, Marjorie Rowe, Eleanor D. Schonberg, Stefanie L. Rodrigues, Austin LaPiana, Robert C. Patio, Trisha Anand, Jana Fisher, Camille Mazurek, Ruoran Guan, Kshitij Wagh, James Theiler, Bette Korber, Dan H. Barouch

**Author notes:** Corresponding author: Dan H. Barouch, M.D., Ph.D., Center for Virology and Vaccine Research, 330 Brookline Avenue, E/CLS-1043, Boston, MA 02115.

## Abstract

The continued evolution of SARS-CoV-2 may lead to evasion of vaccine immunity and natural immunity. A highly mutated Omicron variant BA.2.86 has recently been identified with over 30 amino acid changes in Spike compared with BA.2 and XBB.1.5. As of September 4, 2023, BA.2.86 has been identified in 37 sequences from 10 countries, which is likely an underestimate due to limited surveillance. The ability of BA.2.86 to evade NAbs compared with other currently circulating Omicron variants remains unknown. Our data show that NAb responses to BA.2.86 were lower than to BA.2 but were comparable or slightly higher than to the current circulating recombinant variants XBB.1.5, XBB.1.16, EG.5, EG.5.1, and FL.1.5.1.

---

The continued evolution of SARS-CoV-2 may lead to evasion of vaccine immunity and natural immunity^1-3^. A highly mutated Omicron variant BA.2.86 has recently been identified with over 30 amino acid changes in Spike compared with BA.2 and XBB.1.5 (**Figs. 1A, S1, S2**). As of September 4, 2023, BA.2.86 has been identified in 37 sequences from 10 countries (**Tables S1, S2**), which is likely an underestimate due to limited surveillance. The ability of BA.2.86 to evade NAbs compared with other currently circulating Omicron variants remains unknown.

**Figure 1.**
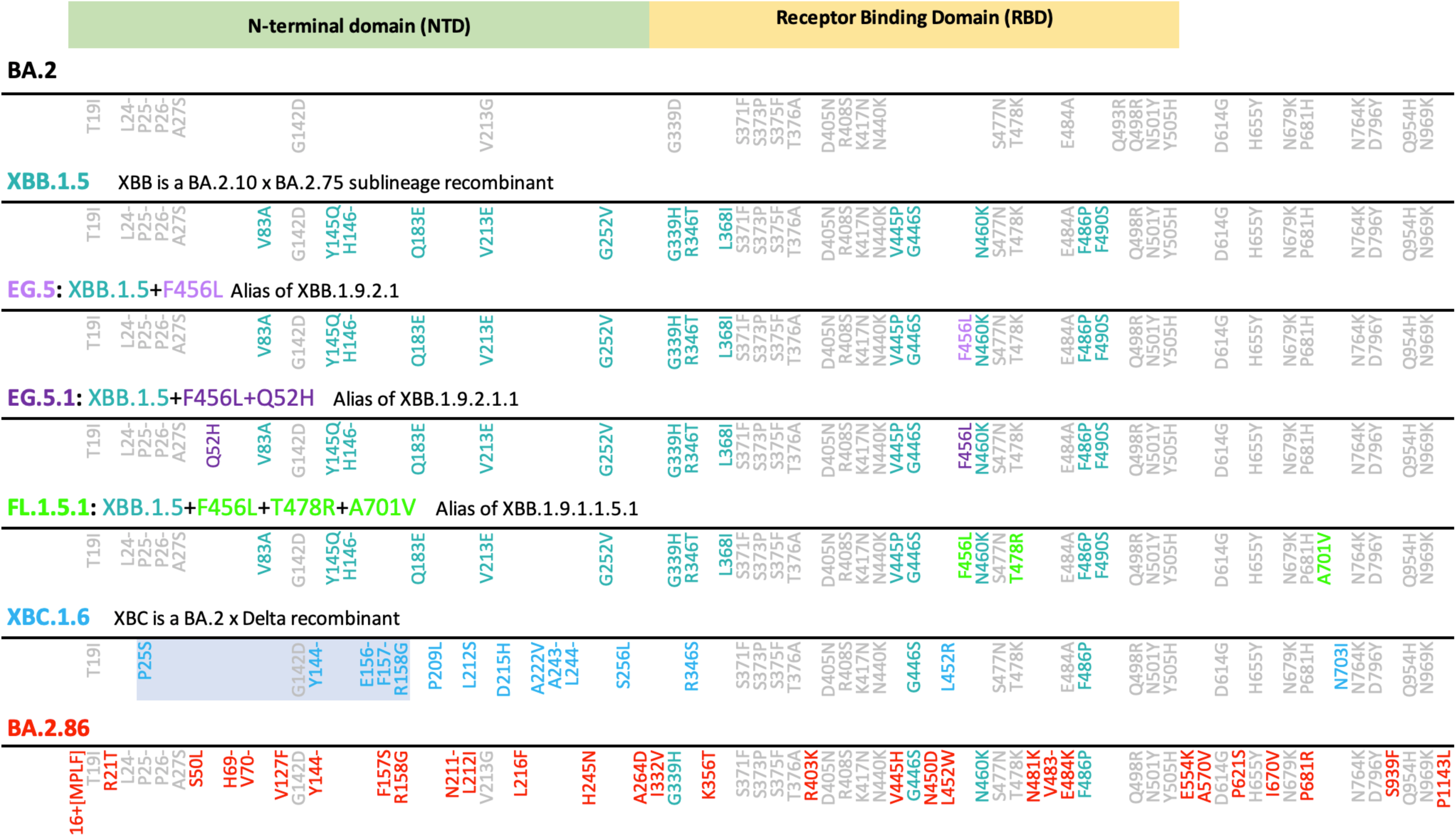
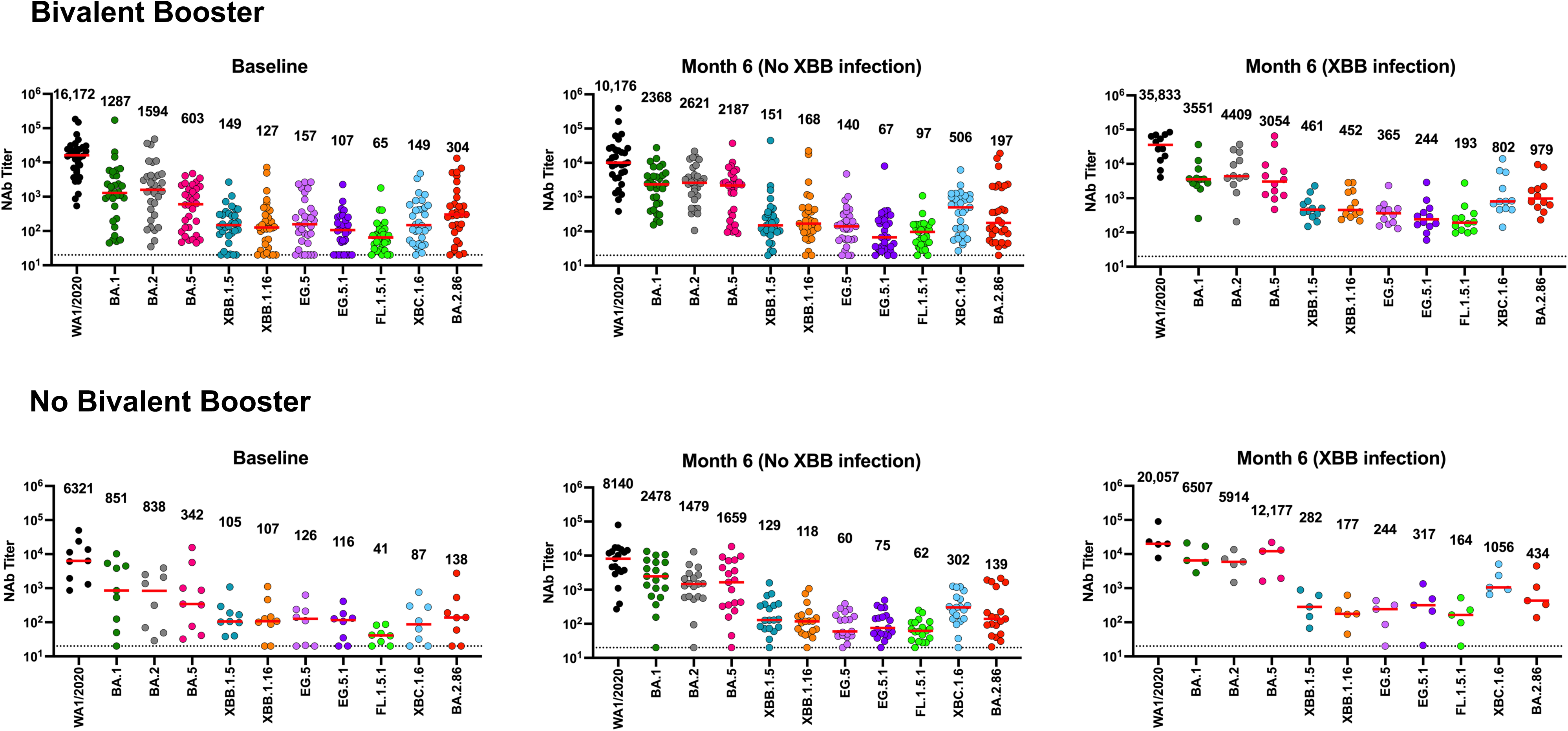
Neutralization escape from SARS-CoV-2 variants. (A) Spike mutations in current circulating SARS-CoV-2 variants. Substitutions in the ancestral BA.2 Omicron lineage relative the Wuhan/WIV04/ reference strain (https://gisaid.org/WIV04/) are shown in grey. Additional substitutions found in XBB.1.5 are highlighted in teal. Additional substitutions in all other study variants relative to these two patterns are indicated for each variants studied. The mutational pattern used matched the consensus form of the expanding lineage, although in BA.2.86 the I670V mutation was present the two earliest sampled intact sequences but has proven to be a rare mutation. The highlighted region in XBC.1.6 is a recombinant fragment from a Delta variant. (B) Neutralizing antibody (NAb) against the WA1/2020, BA.1, BA.2, BA.5, XBB.1.5, XBB.1.16, EG.5, EG.5.1, FL.1.5.1, XBC.1.6, and BA.2.86 variants by luciferase-based pseudovirus neutralization assays at baseline and at 6 months in individuals who received or did not receive the bivalent mRNA booster in fall 2022. Also shown are participants at 6 months who were infected during the XBB.1.5 predominant period. The horizontal red bar reflects median values.

We evaluated NAb responses in 44 individuals who received the bivalent mRNA booster in fall 2022 and in 22 individuals who did not receive the bivalent mRNA booster (**Tables S3, S4**). Participants had a median of 3 COVID-19 vaccine doses prior to the bivalent mRNA boost, and 70-73% had at least one documented SARS-CoV-2 infection. In participants who received the bivalent mRNA boost, baseline NAb responses against WA1/2020, BA.1, BA.2, BA.5, XBB.1.5, XBB.1.16, EG.5, EG.5.1, FL.1.5.1, XBC.1.6, and BA.2.86 were 16,172, 1287, 1594, 603, 149, 127, 157, 107, 65, 149, and 304, respectively (**Fig. 1B**). NAb responses increased at 3 weeks after the boost (**Fig. S3**) and declined largely back to baseline at 6 months in individuals who did not have a documented infection during the XBB.1.5 predominant period, with NAb responses of 10,176, 2368, 2621, 2187, 151, 168, 140, 67, 97, 506, and 197, respectively. At 6 months in individuals who had a documented infection during the XBB.1.5 predominant period, NAb responses were 35,833, 3551, 4409, 3054, 461, 452, 365, 244, 193, 802, and 979, respectively. Participants who did not receive the bivalent boost had comparable NAb responses at 6 months.

Our data show that NAb responses to BA.2.86 were 5-13 fold lower than to BA.2 but were comparable or slightly higher than to XBB.1.5, XBB.1.16, EG.5, EG.5.1, and FL.1.5.1. BA.2.86 likely evolved directly from the less resistant BA.2 variant, rather than from the current highly resistant circulating recombinant variants, which presumably were selected for increased NAb escape following infection with XBB lineage viruses. XBC.1.6 is another highly mutated variant that is a BA.2/Delta recombinant (**Figs. S4, S5**) and similarly shows less NAb escape than XBB.1.5. Our data also show that NAb profiles at 6 months were comparable in participants who did or did not receive the bivalent mRNA boost, consistent with its limited durability^4,5^, and NAb titers increased substantially following XBB infection. It will be important to monitor for potential further evolution or recombination of BA.2.86.

## Correspondence

Correspondence and requests for materials should be addressed to D.H.B.

## Supplementary Methods

### Study Population

A specimen biorepository at Beth Israel Deaconess Medical Center (BIDMC) obtained samples from individuals who received monovalent SARS-CoV-2 vaccines as well as bivalent mRNA boosters. The BIDMC institutional review board approved this study (2020P000361). All participants provided informed consent. This study included 66 individuals who received the bivalent mRNA booster in fall 2022 (N=44) or who did not receive the bivalent mRNA booster (N=22). Participants were followed for 6 months and were excluded if they received immunosuppressive medications.

### Pseudovirus Neutralizing Antibody Assay

Neutralizing antibody (NAb) titers against SARS-CoV-2 variants were determined using pseudo-typed viruses expressing a luciferase reporter gene. In brief, a luciferase reporter plasmid pLenti-CMV Puro-Luc (Addgene), packaging construct psPAX2 (AIDS Resource and Reagent Program), and Spike protein expressing pcDNA3.1-SARS-CoV-2 SΔCT were co-transfected into human embryonic kidney (HEK)293T cells (ATCC CRL_3216) with lipofectamine 2000 (ThermoFisher Scientific). Pseudo-typed viruses of SARS-CoV-2 variants were generated using the Spike protein from WA1/2020 (Wuhan/WIV04/2019, GISAID accession ID: EPI_ISL_402124), Omicron BA.1 (GISAID ID: EPI_ISL_7358094.2), BA.2 (GISAID ID: EPI_ISL_6795834.2), BA.5 (GISAID ID: EPI_ISL_12268495.2), XBB.1.5 (GISAID ID: EPI_ISL_16418320), XBB.1.16 (GISAID ID: EPI_ISL_17646715), EG.5 (GISAID ID: EPI_ISL_17976635), EG.5.1 (GISAID ID: EPI_ISL_18125149), FL.1.5.1 (GISAID ID: EPI_ISL_18126515), XBC.1.6 (GISAID ID: EPI_ISL_17851490), and BA.2.86 (GISAID ID: EPI_ISL_18110065). 48 hours post-transfection, the supernatants containing the pseudo-typed viruses were collected and purified by filtration with 0.45-μm filter. To determine NAb titers in human sera, HEK293T-hACE2 cells were seeded in 96-well tissue culture plates at a density of 2 × 10^4^ cells per well overnight. Three-fold serial dilutions of heat-inactivated serum samples were prepared and mixed with 60 μl of pseudovirus, and incubated at 37 °C for 1 h before adding to HEK293T-hACE2 cells. 48 h later, cells were lysed in Steady-Glo Luciferase Assay (Promega) according to the manufacturer’s instructions. SARS-CoV-2 neutralization titers were defined as the sample dilution at which a 50% reduction (NT50) in relative light units was observed relative to the average of the virus control wells.

### GISAID Data

We gratefully acknowledge the many data contributors to GISAID, the authors and originating laboratories, who share their data, as well as the GISAID staff who facilitate data dissemination. Their work enables an informed response to newly emerging SARS-CoV-2 variants. In particular, we thank the groups that deposited BA.2.86 data. The following acknowledgement link was provided by GISAID for the BA.2.86 data, for the global data sampled in 2023, and for the Australian data and data from the Philippines sampled in 2022 and 2023. All sequences in this dataset are compared relative to hCoV-19/Wuhan/WIV04/2019 (WIV04), the official reference sequence employed by GISAID (EPI_ISL_402124).

#### BA.2.86

EPI_SET_230904un is composed of 37 individual genome sequences. The collection dates range from 2023-07-24 to 2023-08-26; data were collected in 10 countries and territories. All genome sequences and associated metadata in this dataset are published in GISAID’s EpiCoV database. To view the contributors of each individual sequence with details such as accession number, Virus name, Collection date, Originating Lab and Submitting Lab and the list of Authors, see 10.55876/gis8.230904un

#### All 2023 data through August 31, 2023

EPI_SET_230901af is composed of 932,596 individual genome sequences. The collection dates range from 2023-01-01 to 2023-08-30; data were collected in 157 countries and territories. All genome sequences and associated metadata in this dataset are published in GISAID’s EpiCoV database. To view the contributors of each individual sequence with details such as accession number, Virus name, Collection date, Originating Lab and Submitting Lab and the list of Authors, see 10.55876/gis8.230901af

#### All 2022-2023 data through August 31, 2023 from the Philippines

EPI_SET_230902tu is composed of 14,185 individual genome sequences. The collection dates range from 2022-01-01 to 2023-08-09; data were collected in 1 countries and territories. All genome sequences and associated metadata in this dataset are published in GISAID’s EpiCoV database. To view the contributors of each individual sequence with details such as accession number, Virus name, Collection date, Originating Lab and Submitting Lab and the list of Authors, see 10.55876/gis8.230902tu

#### All 2022-2023 data through August 31, 2023 from Australia

EPI_SET_230902sy is composed of 157,103 individual genome sequences. The collection dates range from 2022-01- 01 to 2023-08-22; data were collected in 1 countries and territories. All genome sequences and associated metadata in this dataset are published in GISAID’s EpiCoV database. To view the contributors of each individual sequence with details such as accession number, Virus name, Collection date, Originating Lab and Submitting Lab and the list of Authors, see 10.55876/gis8.230902sy

---

**Figure S1.**
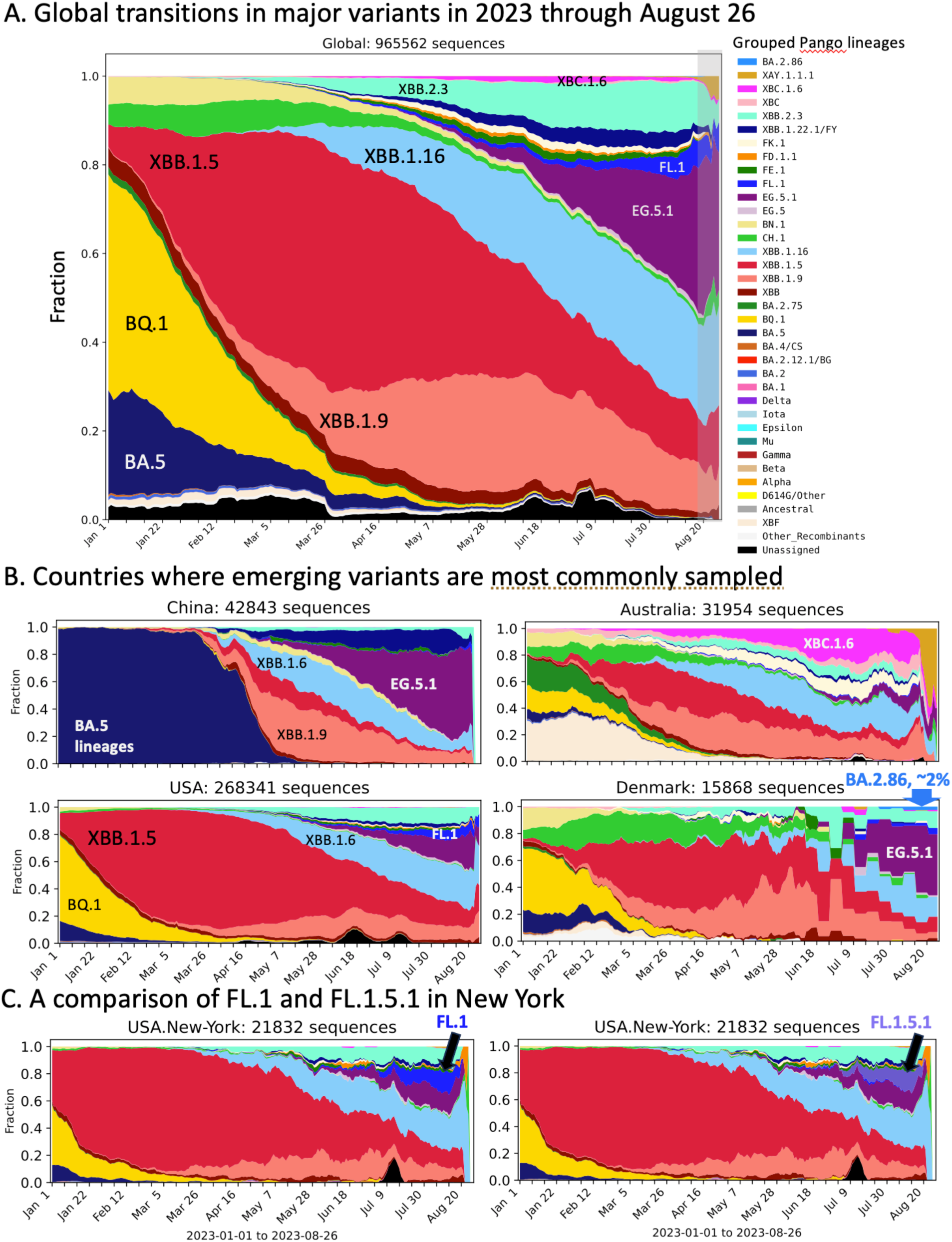
Transitions in SARS-CoV-2 variant lineages in 2023. (A) Global frequencies based on GISAID data. The grey overlay emphasizes that recent sampling is biased due to inevitable time delays between sampling and sequence submissions to GISAID. EG.5.1 sampling is currently increasing the fastest globally. (B) shows the countries were the emergent lineages are most sampled (**Table S1, S2**). EG.5.1 has become the dominant lineage in China, and is increasing everywhere it is sampled. XBC.1 and XBC.1.6 are common in Australia (magenta), but no longer increasing there, while a newly detected variant of an older recombinant lineage, XAY.1.1.1, has recently emerged in Tasmania as a part of the Australian sample (light brown). The recently detected highly divergent variant BA.2.86 has 37 sequences available in GISAID as of this writing, 2023-09-04. It is still too rare to be readily visualized on the global plot in (A), but is beginning to be visually evident in Denmark where it has been sampled 12 times, about 2% of the 542 seqquences submitted by Denmark since the first BA.2.86 was sampled there on 2023-07-24. The FL.1.5.1 sublineage is increasing quickly in the USA, but it has not yet been designated in GISAID, so we so the FL.1 designation instead. As FL.1.5.1 is by far the dominant sublineage among recent FL.1 designated samples they track closely. (C) New York is showing the most rapid increase in FL.1.5.1 sampling frequency at the state level. FL.1.5.1 can be differentiaed from FL.1 by 2 additional mutations in Spike, F456L and T478R. When we track FL.1 designated sequences in New York (blue, left), versus tracking that actual subset of sequences that are defined by FL.1+ F456L +T478R (slate blue, right), it is evident that FL.1.5.1 is the dominant recently expanding form within the FL.1 lineage variants.

**Figure S2.**
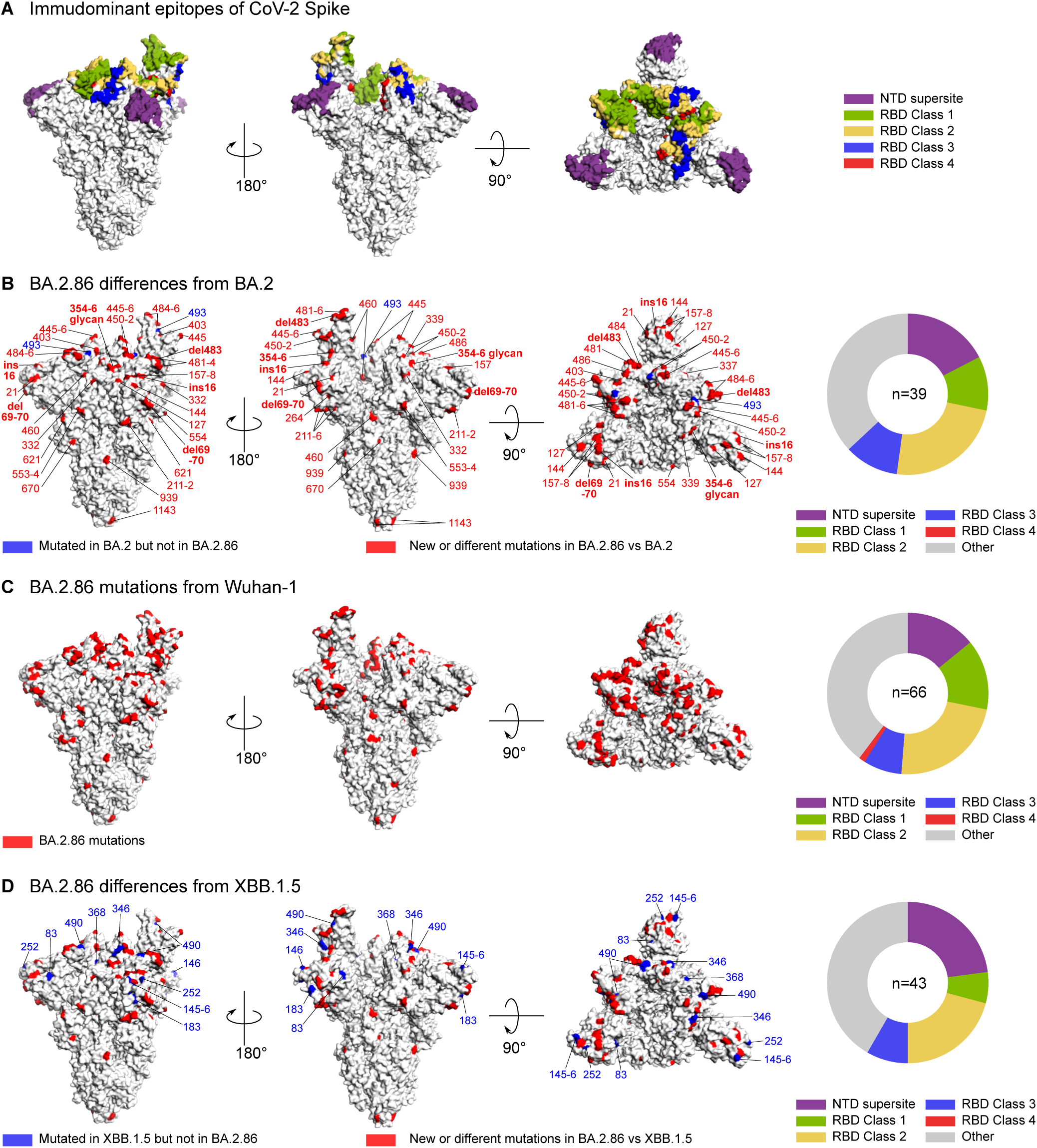
Structural models of BA.2.86 mutations. (A) Structural mapping of key neutralizing antibody epitopes. Spike trimer structure in the one-up configuration from Benton et al. (PMID: 32942285) (PDB: 7A94) is used. RBD class and epitope definitions are from Barnes et al. (PMID: 33045718), and NTD supersite epitope definitions are from Cerutti et al. (PMID: 33789084). Epitopes are color-coded per legend. (B-D) Mutations in BA.2.86 versus BA.2, Wuhan-1, and XBB.1.5. Sites with either new or different mutations in BA.2.86 with respect to the comparator variant are shown in red, and sites that have mutations in the comparator strain but not in BA.2.86 are shown in blue. An N-linked glycosylation site introduced in BA.2.96 at Spike N354 (resulting from a K356T change) and insertions and deletions are in bold as they may be particularly impactful. The pie charts on the right show the fraction of sites with amino acid differences between BA.2.86 and the comparator variant that fall in each of the epitopes from (A); grey indicates fraction of sites with sequence differences that did not fall in any of the epitopes from (A). The total number of sites with sequence differences is indicated in the center, and sites that occur in multiple epitopes are counted in each epitope.

**Figure S3.**
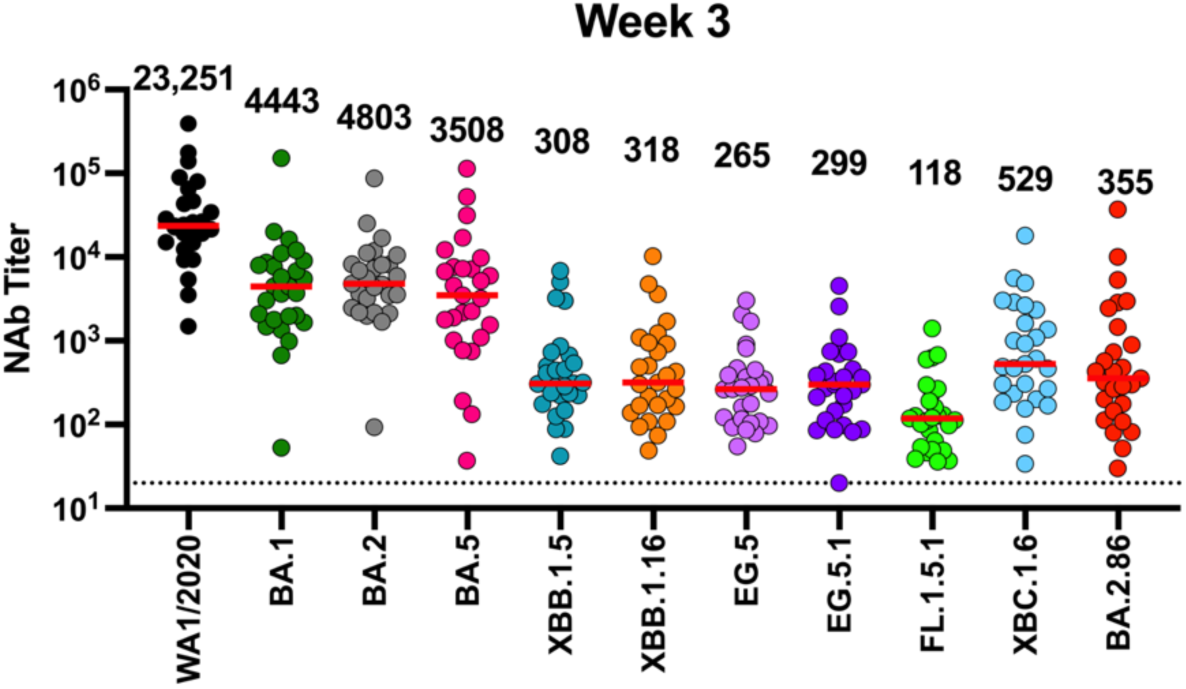
Neutralizing antibody (NAb) against the WA1/2020, BA.1, BA.2, BA.5, XBB.1.5, XBB.1.16, EG.5, EG.5.1, FL.1.5.1, XBC.1.6, and BA.2.86 variants by luciferase-based pseudovirus neutralization assays at week 3 following the bivalent mRNA booster in fall 2022.

**Figure S4.**
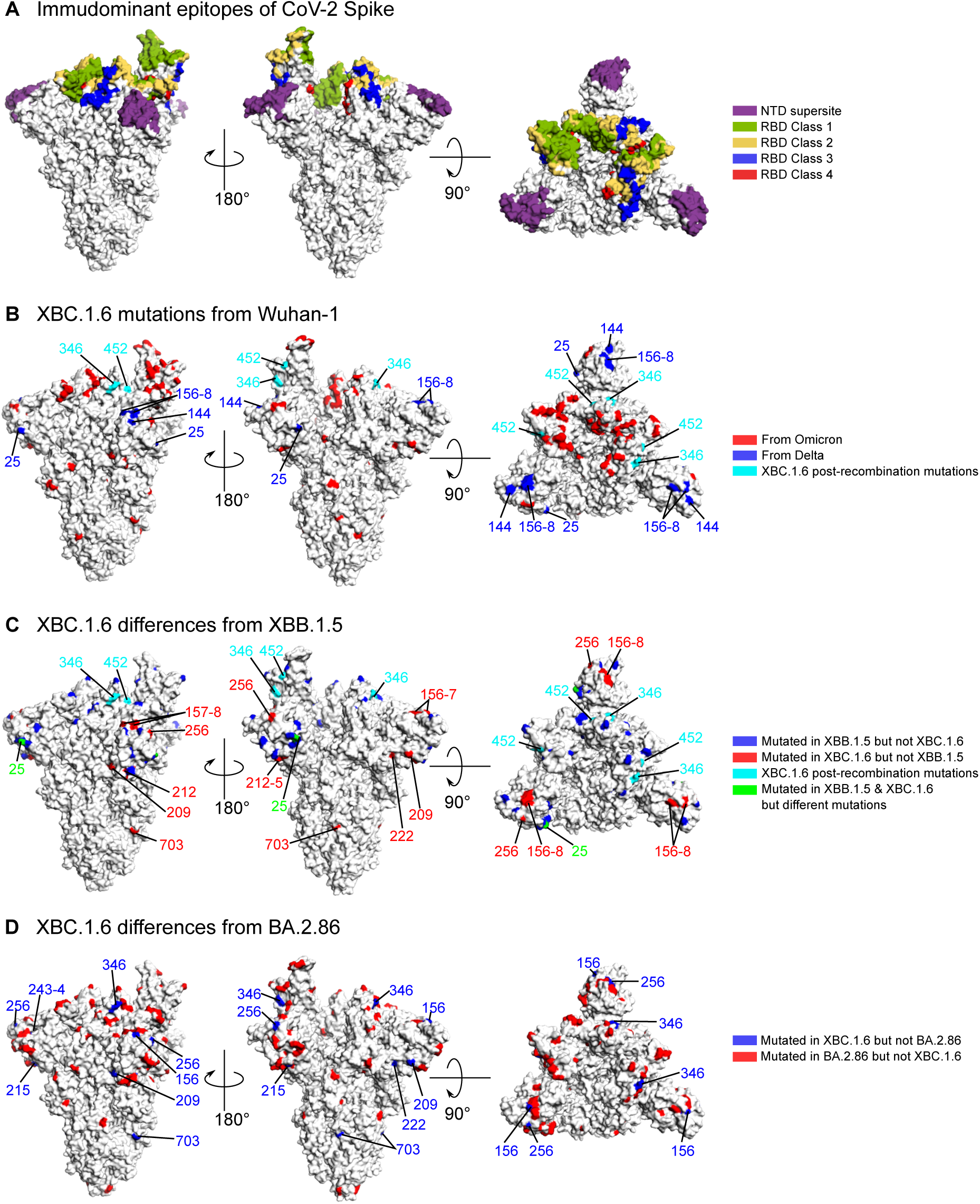
Structural mapping of mutations in XBC.1.6. (A) Same as **Figure S2** panel (A) for reference. (B) XBC.1.6 mutations away from Wuhan-1. Mutations in XBC.1.6 are colored red if they originate from the putative Omicron recombination parent, blue if they originate from the Delta recombination parent, and cyan if they putatively arose after the recombination event. (C) XBC.1.6 differences from XBB.1.5. Sites with mutations in XBB.1.5 but not in XBC.1.6 are shown in blue, and sites with mutations in XBC.1.6 but not in XBB.1.5 are shown in red. Sites 346 and 452 with post-recombination mutations in XBC.1.6 are shown in cyan. The sole site 25 that was mutated in both XBC.1.6 and XBB.1.5 but with different amino acids is shown in green. (D) XBC.1.6 differences from BA.2.86. Sites with new or different mutations in BA.2.86 compared to XBC.1.6 are shown in red and sites with mutations in XBC.1.6 but not in BA.2.86 are shown in blue.

**Figure S5.**
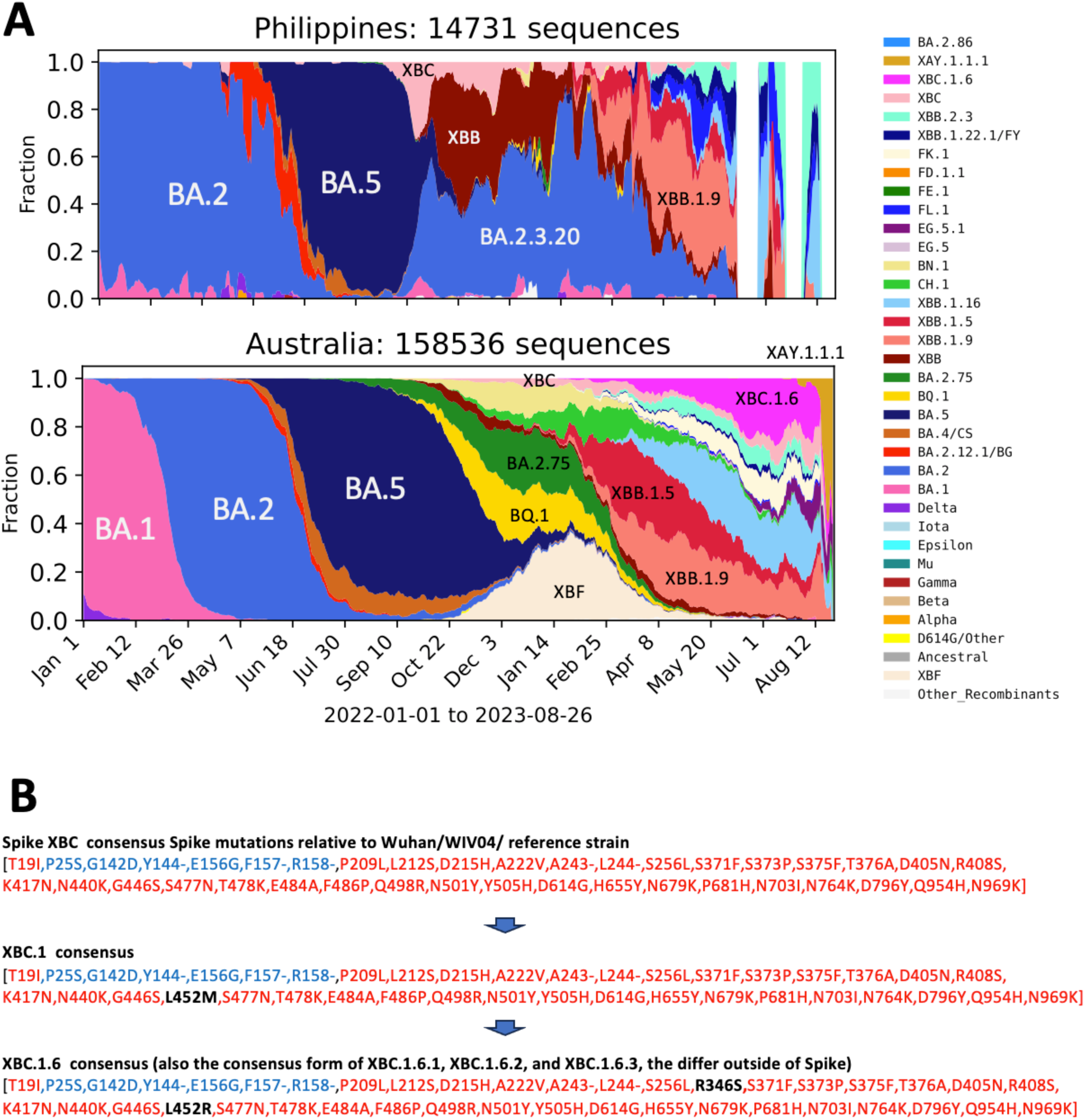
Origin and trajectory of XBC.1.6. A. The Delta-Omicron XBC recombinant first became apparent in the Philippines, where it began to steeply rise in sampling frequency in the fall of 2022. The increase was transient, and XBC and XBC.1 soon was replaced by XBB and BA.2.3.20 lineages. Prior to its decline it had begun to spread internationally, seeding a region spread Australia. The original XBC recombinant form did not expand, but the XBC.1.6 form which had acquired two additional mutations, L452R and R346 began to be increasingly sampled in Australia throughout the spring of 2023, and during this period of expansion XBC.1.6 began to be sampled globally (see **Table S1**). **B.** Exact matches to potential parental variants that gave rise to the XBC recombinant are difficult to find in GISAID, but examples of variants that may be related to parental lineages include the Delta AY.122 variant EPI_ISL_7212429, which exactly matches the amino acid pattern shown in blue, and the Omicron B.1.1.529 variant EPI_ISL_16017283, which is not a perfect match but relative closely resembles the amino acid pattern shown in red.

**Table S1.** Sampling counts and geographic origins of emerging lineages in this study.

| Counts in GISAID, 2023/09/01 | Pango sublineage designations not yet in GISAID but that are captured in the larger grouping in column A* | Number of countries, and counts in 2023 for the three most commonly sampled countries | 2023-01-01 to 2023-09-01 | 2023/07/01 to 2023/09/01 | % sampled in the last 2 months |
| --- | --- | --- | --- | --- | --- |
| EG.5 |  | global: USA (311), Canada (37), China (38) | 1,368 | 619 | 45.20% |
| EG.5.1 | EG.5.1*, HK.*, JG.1, JJ.1, HV.1 | global: China (4,446), USA (3,772), Japan (2,022) | 17,443 | 13,379 | 76.70% |
| XBC.1.6 | XBC.1.6.4 | 15 countries: Australia (1185), New Zealand (31), USA (27) | 1,269 | 274 | 21.60% |
| XBC.1.6.1 | GT.1 | 7 countries: Australia (370), South Korea (4), China (3) | 385 | 38 | 9.90% |
| XBC.1.6.2 |  | 5 countries: 241 Australia (241), Japan (9), USA (6) | 258 | 57 | 22.00% |
| XBC.1.6.3 | HW.* | 15 countries: Australia (491), USA (46) New Zealand (42) | 645 | 186 | 28.80% |
| FL.1 + F456L + T478R** | FL.1.5, FL.1.5.1 | global: USA (1026), Canada (198), Dominican Republic (178) | 1,739 | 1,573 | 90.40% |
| BA.2.86 |  | 10 countries: Denmark (12), Sweden (5), USA (4) | 37 | 37 | 100% |

\*Cornelius Roemer's Pango lineage: https://github.com/cov-lineages/pango-designation/blob/master/lineage_notes.txt

**Table S2.** BA.2.86 variants available in GSAID as of September 4, 2023.

| Virus name | Accession ID | Collection date | Location | Host | Sampling strategy |
| --- | --- | --- | --- | --- | --- |
| hCoV-19/South Africa/NICD-N55967/2023 | EPI_ISL_18125259 | 7/24/23 | Africa / South Africa / Gauteng | Human | Baseline Surveillance |
| hCoV-19/South Africa/NICD-R09343/2023 | EPI_ISL_18168185 | 8/17/23 | Africa / South Africa / Gauteng | Human | Baseline Surveillance |
| hCoV-19/South Africa/NICD-N55999/2023 | EPI_ISL_18125249 | 7/28/23 | Africa / South Africa / Mpumalanga | Human | Baseline Surveillance |
| hCoV-19/Israel/ICH-741198454/2023 | EPI_ISL_18096761 | 7/31/23 | Asia / Israel | Human |  |
| hCoV-19/Israel/SMC-7114868/2023 | EPI_ISL_18213454 | 8/26/23 | Asia / Israel | Human |  |
| hCoV-19/env/Thailand/CU-074/2023 | EPI_ISL_18216958 | 7/28/23 | Asia / Thailand / Bangkok | Environment | Wastewater surveillance |
| hCoV-19/env/Thailand/CU-064/2023 | EPI_ISL_18216959 | 7/28/23 | Asia / Thailand / Bangkok | Environment | Wastewater surveillance |
| hCoV-19/env/Thailand/CU-082/2023 | EPI_ISL_18216960 | 7/28/23 | Asia / Thailand / Bangkok | Environment | Wastewater surveillance |
| hCoV-19/env/Thailand/CU-077/2023 | EPI_ISL_18216961 | 7/28/23 | Asia / Thailand / Bangkok | Environment | Wastewater surveillance |
| hCoV-19/Denmark/DCGC-647676/2023 | EPI_ISL_18097345 | 7/24/23 | Europe / Denmark | Human |  |
| hCoV-19/Denmark/DCGC-647646/2023 | EPI_ISL_18097315 | 7/31/23 | Europe / Denmark | Human |  |
| hCoV-19/Denmark/DCGC-647694/2023 | EPI_ISL_18114953 | 8/7/23 | Europe / Denmark | Human |  |
| hCoV-19/Denmark/DCGC-658505/2023 | EPI_ISL_18159709 | 8/7/23 | Europe / Denmark | Human |  |
| hCoV-19/Denmark/DCGC-658800/2023 | EPI_ISL_18160063 | 8/7/23 | Europe / Denmark | Human |  |
| hCoV-19/Denmark/DCGC-647853/2023 | EPI_ISL_18135401 | 8/14/23 | Europe / Denmark | Human |  |
| hCoV-19/Denmark/DCGC-656489/2023 | EPI_ISL_18157710 | 8/14/23 | Europe / Denmark | Human |  |
| hCoV-19/Denmark/DCGC-657228/2023 | EPI_ISL_18158448 | 8/14/23 | Europe / Denmark | Human |  |
| hCoV-19/Denmark/DCGC-657658/2023 | EPI_ISL_18158884 | 8/14/23 | Europe / Denmark | Human |  |
| hCoV-19/Denmark/DCGC-658990/2023 | EPI_ISL_18210602 | 8/14/23 | Europe / Denmark | Human |  |
| hCoV-19/Denmark/DCGC-657232/2023 | EPI_ISL_18158452 | 8/21/23 | Europe / Denmark | Human |  |
| hCoV-19/Denmark/DCGC-658978/2023 | EPI_ISL_18210550 | 8/21/23 | Europe / Denmark | Human |  |
| hCoV-19/France/COR-HCL-723000998801/2023 | EPI_ISL_18213025 | 8/21/23 | Europe / France / Corse | Human |  |
| hCoV-19/France/GES-IPP17379/2023 | EPI_ISL_18168588 | 8/21/23 | Europe / France / Grand Est | Human | Baseline surveillance |
| hCoV-19/Portugal/PT49343/2023 | EPI_ISL_18142273 | 8/15/23 | Europe / Portugal | Human | Representative |
| hCoV-19/Portugal/PT49347/2023 | EPI_ISL_18142275 | 8/15/23 | Europe / Portugal | Human | Representative |
| hCoV-19/Sweden/AB-000-117-986-1/2023 | EPI_ISL_18147545 | 8/7/23 | Europe / Sweden / Stockholm | Human |  |
| hCoV-19/Sweden/AB-01_SE100_CS101480/2023 | EPI_ISL_18151536 | 8/13/23 | Europe / Sweden / Stockholm | Human |  |
| hCoV-19/Sweden/Y-261733286655/2023 | EPI_ISL_18147561 | 8/18/23 | Europe / Sweden / Vasternorrland | Human |  |
| hCoV-19/Sweden/Y-261733291192/2023 | EPI_ISL_18147559 | 8/19/23 | Europe / Sweden / Vasternorrland | Human |  |
| hCoV-19/Sweden/U-M988749/2023 | EPI_ISL_18168405 | 8/18/23 | Europe / Sweden / Vastmanland | Human |  |
| hCoV-19/England/CLIMB-CM7YMG9C/2023 | EPI_ISL_18209338 | 8/16/23 | Europe / United Kingdom / England | Human |  |
| hCoV-19/England/GSTT-YYBYBN4/2023 | EPI_ISL_18111770 | 8/13/23 | Europe / United Kingdom / England / London | Human |  |
| hCoV-19/Canada/BC-BCCDC-641728/2023 | EPI_ISL_18164467 | 8/23 | North America / Canada / British Columbia | Human | Baseline surveillance |
| hCoV-19/USA/MI-UM-10052670540/2023 | EPI_ISL_18110065 | 8/3/23 | North America / USA / Michigan | Human | Baseline surveillance |
| hCoV-19/USA/OH-ODH-SC3032044/2023 | EPI_ISL_18138566 | 7/29/23 | North America / USA / Ohio / Cuyahoga | Human | Baseline surveillance |
| hCoV-19/USA/TX-HMH-M-132251/2023 | EPI_ISL_18164159 | 8/22/23 | North America / USA / Texas / Houston | Human |  |
| hCoV-19/USA/VA-GBW-H20-330-6734/2023 | EPI_ISL_18121060 | 8/10/23 | North America / USA / Virginia / Loudoun County | Human | Travel history: Asia / Japan |

**Table S3.**
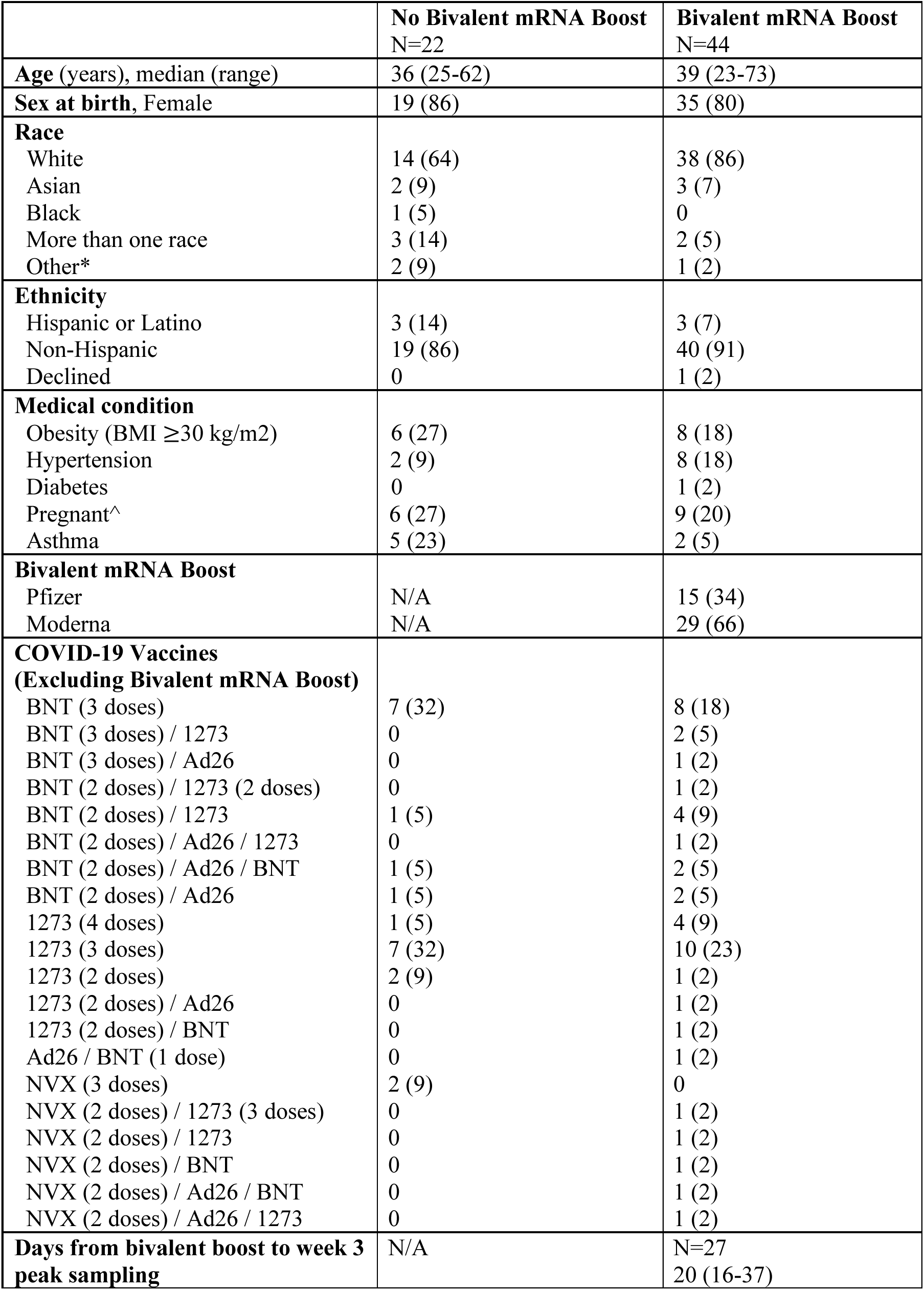
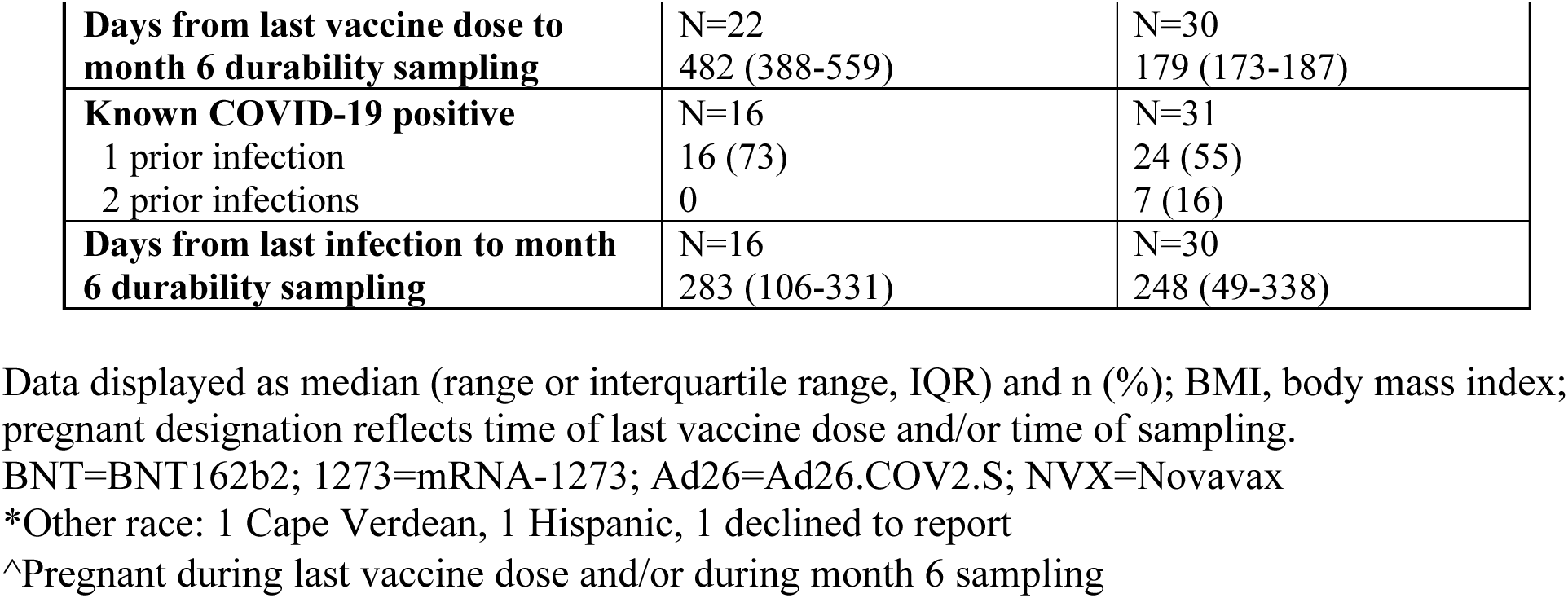
Study population.

|  | **No Bivalent mRNA Boost**<br>N=22 | **Bivalent mRNA Boost**<br>N=44 |
| --- | --- | --- |
| **Age** (years), median (range) | 36 (25-62) | 39 (23-73) |
| **Sex at birth**, Female | 19 (86) | 35 (80) |
| **Race** |  |  |
| White | 14 (64) | 38 (86) |
| Asian | 2 (9) | 3 (7) |
| Black | 1 (5) | 0 |
| More than one race | 3 (14) | 2 (5) |
| Other* | 2 (9) | 1 (2) |
| **Ethnicity** |  |  |
| Hispanic or Latino | 3 (14) | 3 (7) |
| Non-Hispanic | 19 (86) | 40 (91) |
| Declined | 0 | 1 (2) |
| **Medical condition** |  |  |
| Obesity (BMI $\geq 30$ kg/m<sup>2</sup>) | 6 (27) | 8 (18) |
| Hypertension | 2 (9) | 8 (18) |
| Diabetes | 0 | 1 (2) |
| Pregnant^ | 6 (27) | 9 (20) |
| Asthma | 5 (23) | 2 (5) |
| **Bivalent mRNA Boost** |  |  |
| Pfizer | N/A | 15 (34) |
| Moderna | N/A | 29 (66) |
| **COVID-19 Vaccines**<br>**(Excluding Bivalent mRNA Boost)** |  |  |
| BNT (3 doses) | 7 (32) | 8 (18) |
| BNT (3 doses) / 1273 | 0 | 2 (5) |
| BNT (3 doses) / Ad26 | 0 | 1 (2) |
| BNT (2 doses) / 1273 (2 doses) | 0 | 1 (2) |
| BNT (2 doses) / 1273 | 1 (5) | 4 (9) |
| BNT (2 doses) / Ad26 / 1273 | 0 | 1 (2) |
| BNT (2 doses) / Ad26 / BNT | 1 (5) | 2 (5) |
| BNT (2 doses) / Ad26 | 1 (5) | 2 (5) |
| 1273 (4 doses) | 1 (5) | 4 (9) |
| 1273 (3 doses) | 7 (32) | 10 (23) |
| 1273 (2 doses) | 2 (9) | 1 (2) |
| 1273 (2 doses) / Ad26 | 0 | 1 (2) |
| 1273 (2 doses) / BNT | 0 | 1 (2) |
| Ad26 / BNT (1 dose) | 0 | 1 (2) |
| NVX (3 doses) | 2 (9) | 0 |
| NVX (2 doses) / 1273 (3 doses) | 0 | 1 (2) |
| NVX (2 doses) / 1273 | 0 | 1 (2) |
| NVX (2 doses) / BNT | 0 | 1 (2) |
| NVX (2 doses) / Ad26 / BNT | 0 | 1 (2) |
| NVX (2 doses) / Ad26 / 1273 | 0 | 1 (2) |
| **Days from bivalent boost to week 3 peak sampling** | N/A | N=27<br>20 (16-37) |
| **Days from last vaccine dose to month 6 durability sampling** | N=22<br>482 (388-559) | N=30<br>179 (173-187) |
| **Known COVID-19 positive** | N=16 | N=31 |
| 1 prior infection | 16 (73) | 24 (55) |
| 2 prior infections | 0 | 7 (16) |
| **Days from last infection to month 6 durability sampling** | N=16<br>283 (106-331) | N=30<br>248 (49-338) |

Data displayed as median (range or interquartile range, IQR) and n (%); BMI, body mass index; pregnant designation reflects time of last vaccine dose and/or time of sampling.
BNT=BNT162b2; 1273=mRNA-1273; Ad26=Ad26.COV2.S; NVX=Novavax
\*Other race: 1 Cape Verdean, 1 Hispanic, 1 declined to report
^Pregnant during last vaccine dose and/or during month 6 sampling

**Table S4.** Individual participant level data.

|  | **Age** | **Sex** | **Race** | **Ethnicity** | **Medical history** | **Bivalent boost** | **Vaccine history** | **Bivalent boost to peak (d)** | **Last dose to 6 mo sample (d)** | **# Prior Infections** | **Last infection to 6 mo sample (d)** |
| --- | --- | --- | --- | --- | --- | --- | --- | --- | --- | --- | --- |
| Bival-1 | 32 | M | White | Not H/L |  | Pfizer | NVX/NVX/1273 | N/A | 186 | 0 | N/A |
| Bival-2 | 43 | F | White | Not H/L |  | Moderna | BNT/BNT/BNT | 28 | 182 | 0 | N/A |
| Bival-3 | 48 | M | White | Not H/L | Asthma | Moderna | 1273/1273/BNT | 62 | 179 | 2 | 43 |
| Bival-4 | 37 | F | White | Not H/L | Obesity | Moderna | NVX/NVX/Ad26/1273 | N/A | 190 | 1 | 83 |
| Bival-5 | 68 | M | White | Not H/L | Obesity, HTN | Moderna | 1273/1273/1273/1273 | N/A | 184 | 1 | 17 |
| Bival-6 | 37 | F | White | Not H/L |  | Moderna | 1273/1273/1273 | 39 | 174 | 0 | N/A |
| Bival-7 | 38 | F | White | H/L |  | Moderna | BNT/BNT/BNT | N/A | 209 | 1 | 56 |
| Bival-8 | 32 | F | White | Not H/L | Pregnancy | Moderna | BNT/BNT/BNT | 25 | 196 | 1 | 306 |
| Bival-9 | 52 | M | White | Not H/L |  | Moderna | 1273/1273/1273 | N/A | 204 | 2 | 259 |
| Bival-10 | 37 | F | Asian | Not H/L |  | Pfizer | BNT/BNT/BNT/1273 | 15 | 189 | 0 | N/A |
| Bival-11 | 27 | M | Multiple races | H/L |  | Moderna | 1273/1273/1273 | N/A | 195 | 1 | 333 |
| Bival-12 | 36 | F | White | Not H/L | Obesity, Pregnancy | Moderna | BNT/BNT/BNT | N/A | 167 | 0 | N/A |
| Bival-13 | 29 | F | Asian | Not H/L | Pregnancy, Asthma | Moderna | BNT/BNT/1273 | N/A | 185 | 1 | 125 |
| Bival-14 | 36 | F | White | Not H/L |  | Moderna | BNT/BNT/1273 | 37 | 172 | 1 | 155 |
| Bival-15 | 49 | F | White | Not H/L |  | Moderna | 1273/1273/1273 | N/A | 184 | 1 | 13 |
| Bival-16 | 70 | F | White | Not H/L |  | Pfizer | BNT/BNT/Ad26 | 22 | 166 | 0 | N/A |
| Bival-17 | 30 | F | Multiple races | H/L |  | Pfizer | BNT/BNT/BNT | 32 | 174 | 0 | N/A |
| Bival-18 | 46 | F | White | Not H/L |  | Pfizer | BNT/BNT/BNT | 20 | 193 | 0 | N/A |
| Bival-19 | 36 | F | White | Not H/L | HTN, Pregnancy, Asthma | Moderna | BNT/BNT/1273 | 49 | 178 | 1 | 410 |
| Bival-20 | 62 | F | White | Not H/L | HTN, Asthma | Pfizer | BNT/BNT/Ad26 | 30 | 149 | 2 | 14 |
| Bival-21 | 34 | F | White | Not H/L | Pregnancy | Moderna | 1273/1273 | N/A | 166 | 2 | 25 |
| Bival-22 | 66 | F | White | Not H/L | Obesity,<br>HTN | Moderna | BNT/BNT/BNT | 16 | 191 | 0 | N/A |
| Bival-23 | 43 | M | White | Not H/L |  | Pfizer | BNT/BNT/1273 | 15 | 183 | 1 | 466 |
| Bival-24 | 66 | F | White | Not H/L |  | Moderna | 1273/1273/1273/1273 | 72 | 191 | 0 | N/A |
| Bival-25 | 37 | F | White | Not H/L |  | Pfizer | 1273/1273/1273 | 15 | 174 | 0 | N/A |
| Bival-26 | 51 | F | White | Not H/L |  | Pfizer | BNT/BNT/BNT | 65 | 177 | 1 | 1093 |
| Bival-27 | 40 | F | White | Not H/L |  | Pfizer | BNT/BNT/Ad26/1273 | N/A | 173 | 1 | 242 |
| Bival-28 | 40 | F | White | Not H/L |  | Moderna | BNT/BNT/BNT | 33 | 178 | 1 | 305 |
| Bival-29 | 73 | F | White | Not H/L |  | Moderna | 1273/1273/1273/1273 | 20 | 186 | 1 | 332 |
| Bival-30 | 39 | F | White | Not H/L | Obesity,<br>HTN | Pfizer | NVX/NVX/BNT | N/A | 193 | 1 | 55 |
| Bival-31 | 34 | F | White | Not H/L | Pregnancy | Moderna | 1273/1273/1273 | 19 | 175 | 2 | 314 |
| Bival-32 | 23 | F | White | Not H/L | Obesity | Pfizer | 1273/1273/Ad26 | 14 | 183 | 1 | 358 |
| Bival-33 | 41 | M | White | Not H/L |  | Moderna | NVX/NVX/1273/1273/<br>1273 | N/A | 182 | 1 | 365 |
| Bival-34 | 32 | F | White | Not H/L | HTN,<br>Pregnancy | Moderna | BNT/BNT/BNT | 18 | 179 | 1 | 301 |
| Bival-35 | 35 | F | White | Not H/L |  | Moderna | 1273/1273/1273 | 19 | 172 | 0 | N/A |
| Bival-36 | 72 | M | White | Not H/L | HTN,<br>Asthma | Pfizer | BNT/BNT/Ad26 | 55 | 173 | 0 | N/A |
| Bival-37 | 25 | F | White | Not H/L |  | Pfizer | BNT/BNT/Ad26 | 14 | 187 | 1 | 253 |
| Bival-38 | 52 | F | White | Not H/L |  | Moderna | NVX/NVX/Ad26 | N/A | 173 | 1 | 18 |
| Bival-39 | 43 | F | White | Not H/L |  | Pfizer | 1273/1273/1273 | 20 | 167 | 1 | 51 |
| Bival-40 | 31 | F | White | Not H/L | Pregnancy,<br>Asthma | Moderna | 1273/1273/1273 | 17 | 171 | 1 | 886 |
| Bival-41 | 31 | F | White | Not H/L | Pregnancy | Moderna | 1273/1273/1273 | N/A | 167 | 2 | 105 |
| Bival-42 | 40 | F | Declined | Declined | Obesity,<br>HTN,<br>Diabetes | Moderna | 1273/1273/1273/1273 | N/A | 174 | 1 | 14 |
| Bival-43 | 52 | M | White | Not H/L |  | Moderna | BNT/BNT/1273/1273 | N/A | 184 | 1 | 352 |
| Bival-44 | 27 | F | Asian | Not H/L |  | Moderna | Ad26/BNT | 12 | N/A | 2 | 90 |
| NoBival-1 | 25 | F | Asian | Not H/L |  | N/A | BNT/BNT/Ad26 | N/A | 551 | 0 | N/A |
| NoBival-2 | 50 | F | Multiple races | H/L | Obesity, Asthma | N/A | BNT/BNT/Ad26/BNT | N/A | 196 | 1 | 280 |
| NoBival-3 | 35 | F | White | Not H/L |  | N/A | BNT/BNT/BNT | N/A | 559 | 1 | 443 |
| NoBival-4 | 37 | F | White | Not H/L |  | N/A | 1273/1273/1273 | N/A | 476 | 0 | N/A |
| NoBival-5 | 28 | F | White | Not H/L |  | N/A | 1273/1273/1273 | N/A | 499 | 1 | 186 |
| NoBival-6 | 25 | F | White | Not H/L |  | N/A | 1273/1273/1273 | N/A | 490 | 0 | N/A |
| NoBival-7 | 62 | F | White | Not H/L |  | N/A | 1273/1273 | N/A | 807 | 1 | 285 |
| NoBival-8 | 30 | F | Cape Verdean | Not H/L |  | N/A | BNT/BNT/BNT | N/A | 238 | 1 | 490 |
| NoBival-9 | 27 | F | White | H/L | Pregnancy, Asthma | N/A | 1273/1273/1273 | N/A | 458 | 0 | N/A |
| NoBival-10 | 56 | F | White | Not H/L |  | N/A | BNT/BNT/BNT | N/A | 386 | 1 | 316 |
| NoBival-11 | 31 | F | White | Not H/L | Asthma | N/A | NVX/NVX/NVX | N/A | 419 | 1 | 217 |
| NoBival-12 | 37 | M | Asian | Not H/L |  | N/A | BNT/BNT/BNT | N/A | 796 | 1 | 79 |
| NoBival-13 | 29 | F | Black | Not H/L |  | N/A | 1273/1273/1273 | N/A | 460 | 1 | 336 |
| NoBival-14 | 38 | F | White | Not H/L | Obesity | N/A | 1273/1273/1273 | N/A | 518 | 1 | 454 |
| NoBival-15 | 36 | M | White | Not H/L | Asthma | N/A | NVX/NVX/NVX | N/A | 385 | 1 | 30 |
| NoBival-16 | 38 | M | White | Not H/L | Obesity, HTN | N/A | 1273/1273 | N/A | 719 | 1 | 37 |
| NoBival-17 | 58 | F | Multiple races | Not H/L | Obesity, HTN | N/A | BNT/BNT/BNT | N/A | 388 | 1 | 78 |
| NoBival-18 | 34 | F | White | Not H/L | Pregnancy | N/A | 1273/1273/1273/1273 | N/A | 129 | 1 | 202 |
| NoBival-19 | 31 | F | White | Not H/L | Obesity, Pregnancy | N/A | BNT/BNT/BNT | N/A | 570 | 0 | N/A |
| NoBival-20 | 29 | F | Hispanic | H/L | Pregnancy, Asthma | N/A | 1273/1273/1273 | N/A | 560 | 1 | 293 |
| NoBival-21 | 39 | F | White | Not H/L | Pregnancy | N/A | BNT/BNT/BNT | N/A | 488 | 1 | 306 |
| NoBival-22 | 37 | F | Multiple races | Not H/L | Obesity, Pregnancy | N/A | BNT/BNT/1273 | N/A | 461 | 0 | N/A |

Age reported in years at time of last sampling. D=day; F=female; M=male; H/L=Hispanic/Latino; HTN=hypertension; N/A= not applicable as no sample collected; BNT=BNT162b2; 1273=mRNA-1273; Ad26=Ad26.COV2.S; NVX= Novavax COVID-19 Vaccine

